# fNIRS Dataset During Complex Scene Analysis

**DOI:** 10.1101/2024.01.23.576715

**Authors:** Matthew Ning, Sudan Duwadi, Meryem A. Yücel, Alexander Von Lühmann, David A. Boas, Kamal Sen

## Abstract

When analyzing complex scenes, humans often focus their attention on an object at a particular spatial location. The ability to decode the attended spatial location would facilitate brain computer interfaces for complex scene analysis (CSA). Here, we investigated capability of functional near-infrared spectroscopy (fNIRS) to decode audio-visual spatial attention in the presence of competing stimuli from multiple locations. We targeted dorsal frontoparietal network including frontal eye field (FEF) and intra-parietal sulcus (IPS) as well as superior temporal gyrus/planum temporal (STG/PT). They all were shown in previous functional magnetic resonance imaging (fMRI) studies to be activated by auditory, visual, or audio-visual spatial tasks. To date, fNIRS has not been applied to decode auditory and visual-spatial attention during CSA, and thus, no such dataset exists yet. This report provides an open-access fNIRS dataset that can be used to develop, test, and compare machine learning algorithms for classifying attended locations based on the fNIRS signals on a single trial basis.

## INTRODUCTION

The human brain is an astonishingly powerful computational device, capable of feats yet to be matched by machines. One impressive example is the brain’s ability to selectively attend to specific objects in a complex scene with multiple objects. For example, at a crowded cocktail party, we can look at a friend and hear what they are saying in the midst of other speakers, music, and background noise. Such multisensory filtering allows us to select and process important objects in a complex scene, a process known as Complex Scene Analysis (CSA) (Cherry, 2005; Haykin and Chen, 2005; McDermott, 2009). In stark contrast, millions of humans worldwide with disorders such as ADHD (Mihali et al., 2018; Fu et al., 2022), autism (Marco et al., 2011; *Diagnostic and statistical manual of mental disorders: DSM-5*_TM_, 5th ed, 2013), and hearing losses (Marrone, Mason and Kidd, 2008) find such complex scenes confusing, overwhelming and debilitating. Thus, brain-computer interfaces (BCIs) and assistive devices for CSA have the potential to improve the quality of life for many humans.

CSA can be broken down into several distinct components: determining the spatial location of a target stimulus, segregating the target stimulus from the competing stimuli in the scene, and reconstructing the target stimulus from the mixture forming a perceptual object (Bizley, Maddox and Lee, 2016). Recently, we proposed a brain-inspired algorithm for auditory scene analysis based on a model of cortical neurons (Maddox et al., 2012; Dong, Colburn and Sen, 2016; Chou et al., 2022). This algorithm has the potential to be applied in assistive devices for CSA. However, a critical piece of information required by the algorithm is the spatial location of the target stimulus. Thus, a portable and non-invasive technology that can decode the spatial location of the attended target stimulus during CSA would greatly facilitate the development of BCIs and assistive devices for CSA. In addition, a technology that provides insights into specific brain regions that play a significant role in decoding the attended location has the potential to advance our understanding of fundamental brain mechanisms underlying CSA in both normal and impaired humans.

Functional near-infrared spectroscopy (fNIRS) is a non-invasive neuroimaging technique that measures changes in oxygenated (HbO2) and deoxygenated hemoglobin (HbR) in the cerebral cortex (Chance et al., 1993; Girouard and Iadecola, 2006). Due to its portability and low cost, fNIRS has been used in Brain-Computer Interface (BCI) applications (Naseer and Hong, 2015). Previously, fNIRS has been applied to various aspects of auditory science such as classifying different sound categories (Hong and Santosa, 2016), identifying spatial locations of noise stimuli (Tian et al., 2021), characterizing hemodynamic responses to varying auditory stimuli (Pollonini et al., 2014; Steinmetzger et al., 2020; Luke et al., 2021), and investigating informational masking (Zhang, Mary Ying and Ihlefeld, 2018; Zhang, Alamatsaz and Ihlefeld, 2021). However, to date, fNIRS has not been applied to decode auditory and visual-spatial attention during CSA, and thus, no such dataset exists yet. This report provides an open-access fNIRS dataset that can be used to develop, test, and compare machine learning algorithms for classifying attended locations based on the fNIRS signals on a single trial basis.

Here, we collected brain signals with fNIRS during the presentation of audio-visual stimuli in the presence of competing stimuli from multiple locations in order to mimic complex natural scenes. We targeted the dorsal frontoparietal network including frontal eye field (FEF) and intra-parietal sulcus (IPS) as well as superior temporal gyrus/planum temporal (STG/PT), which were shown to be activated by auditory, visual, or audio-visual spatial tasks in fMRI (Shomstein and Yantis, 2006; Deouell et al., 2007; Wu et al., 2007; Smith et al., 2010; W et al., 2011; Michalka et al., 2015, 2016) and simultaneous magnetoencephalography (MEG)/electroencephalogram (EEG) studies (Larson and Lee, 2013). We also recorded task performance by asking the subjects to identify the content of audio-visual stimuli that they were asked to attend to.

Our sample preliminary analysis using Linear Discriminant Analysis (LDA) with Ledoit and Wolf’s covariance matrix estimator (Ledoit and Wolf, 2004) shows robust decoding of attended spatial location for more than half of our subjects. Analysis code for two-class classification performed in this paper is publicly available on GitHub at https://github.com/NSNC-Lab/fNIRS_DecodingSpatialAttention.

## METHODS

### Participants and Demographics

Twelve adults with normal hearing (age 19-48, 5 males and 7 females) were recruited for this study in accordance with the Institutional Review Board of Boston University. A COVID-19 protocol was developed and strictly adhered to. Participants were screened to exclude those with neurological and psychiatric disorders. Participants were briefed and consented before partaking in this study and were compensated for their time.

### Experimental Paradigm

Participants were seated in front of 3 monitors, located at locations equidistant from the subject: center or 0°, 45° to the left, and 45° to the right. The subjects were provided with chin rest throughout the experiment to discourage head movements. The subjects were asked to refrain from movements during the tasks except for answering the questions at the end of each trial. The auditory stimuli were delivered via an earphone (ER-1 Etymotic Research Inc.) with ear tips (E-A-RLink 3A Insert Eartips), and corresponding videos were displayed on the monitors. The videos used in the experiment are from AVSpeech, a publicly available dataset (Ephrat et al. 2018, https://looking-to-listen.github.io/avspeech/index.html). For each trial, a 2-second long audio-visual cue was delivered randomly at one of the 3 locations in the form of a white cross against a black background and a 2 kHz pure tone linearly ramped in the first 0.5 s. The subjects were instructed to listen to the cue and pay attention to the speaker at the cued location. The cue was followed by 3 videos, one for each location, one of which was the target speaker, and the remaining two were the maskers. The stimuli were followed by two multiple-choice questions on the center monitor, each question containing 5 possible choices. The first question was related to face identification, and the second was related to transcript identification. In the face identification task, the subjects were presented with five different faces and were tasked with correctly identifying the face of the target speaker shown in the video. In the transcript identification task, the subjects were presented with five different transcripts and were tasked with correctly identifying the words spoken by the target speaker in the video. A trial was counted as correct if the participant correctly identified both the face and the words spoken by the target speaker. Upon the completion of two questions, a blank black screen of jittered duration with uniform distribution between 14 and 16 seconds appeared. Thereafter, an instruction to press the space bar to begin the next trial was displayed on the center monitor. While the audio-visual cue and the video clip lasted 2 and 3 seconds respectively, the subjects had 20 seconds in total to answer both multiple-choice questions, but they could move on to the next trial by pressing the space bar immediately after they finished answering.

### Data Acquisition

fNIRS data were collected using continuous wave fNIRS (CW6, Techen System) using 690nm and 830nm wavelengths, with a 50 Hz sampling rate. Multiple channels were supported with frequency multiplexing. The fNIRS recording software was synced to the stimulus presentation with the Neurospec MMB70-817 triggerbox (Neurospec AG, Switzerland). 56cm Landmark Cap (EasyCap, Herrsching, Germany) was used for all the subjects.

### Measurements

#### fNIRS probe design

The probe (optode array) was designed in publicly available AtlasViewer software (https://github.com/BUNPC/AtlasViewer) (Aasted et al., 2015). The probe contains 12 sources, 17 long-separation detectors, and 6 short-separation detectors (SS), for a total of 30 long-separation channels (with an exception for one subject who had 14 sources, 19 long-separation detectors, and 8 short-separation detectors, for a total of 34 long-separation channels). The long-separation detectors were placed 30 mm from the sources whereas SS detectors were placed 8 mm from the sources. The probe covered the dorsal frontoparietal network including the frontal eye field (FEF) and intra-parietal sulcus (IPS) as well as the superior temporal gyrus/planum temporal (STG/PT). Figure 1A and 1B shows the sensitivity profile for the fNIRS probe geometry.

**Figure 1:**
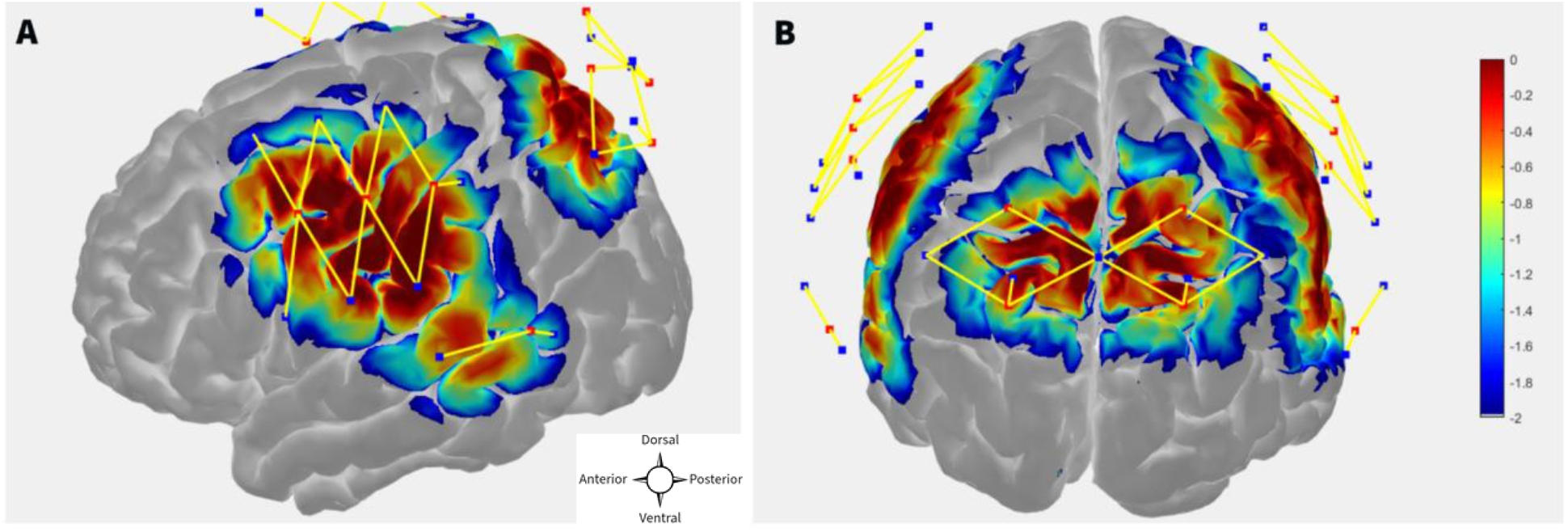
**A, B** fNIRS probe design with 12 sources, 17 regular separation detectors, 6 SS detectors. The sensitivity profile for the probe geometry is shown. Sources and detectors are denoted by red and blue square boxes respectively. The color scale represents the relative sensitivity in log10 units. (Aasted et al., 2015). The longer yellow lines represent regular channels, and the shorter yellow lines represent short separation channels.

### Data structure and format

The fNIRS dataset consists of 90 trials for 11 subjects and 180 trials for 1 subject. The behavioral dataset consists of recorded responses of the subject during each trial and actual answers to the questions asked to the subjects during each trial. fNIRS dataset is in ‘.snirf’ format, and the behavioral dataset is in ‘.mat’ format. Brain Imaging Data Structure (BIDS) compliant fNIRS dataset can be found on OpenNeuro (doi:10.18112/openneuro.ds004830.v1.0.0). The behavioral dataset and the data structure details for both fNIRS and behavioral data can be found on derivatives folder under ‘Experimental and Behavioral Dataset description.docx’.

## PRELIMINARY ANALYSIS AND DATA QUALITY ASSESSMENT

Preliminary fNIRS data preprocessing and analysis were done using publicly available Homer3 software (https://github.com/BUNPC/Homer3) (Huppert et al., 2009). Version of the Homer software used for preliminary analysis is included in GitHub along with custom scripts. Baseline classification analysis using LDA was done using custom code provided at https://github.com/NSNC-Lab/fNIRS_DecodingSpatialAttention.

### Signal-to-noise ratio (SNR)

We excluded subjects that had at least 20 channels pruned using a cutoff of SNR = 1.5, where SNR was estimated as the mean divided by the standard deviation of raw intensity of the fNIRS signal.

### fNIRS pre-processing

Raw light intensities were converted to optical densities using the mean signal as the arbitrary reference point. Motion artifacts in optical density (OD) were identified and corrected with targeted Principal Component Analysis (PCA) before applying criteria for automatic rejection of stimulus trials (Yücel et al., 2014). OD signals were band-pass filtered between 0.01 Hz and 0.5 Hz with a 3rd order zero-phase Butterworth filter. The filtered OD was converted to chromophore concentration changes. The differential path length factor was held fixed at 1, and the concentration unit is in Molar*mm (Scholkmann and Wolf, 2013). Finally, systemic physiological signal clutter was regressed out using a GLM with short-separation channels modeling the signal clutter (Gagnon et al., 2014; von Lühmann et al., 2020). Each regular channel was assigned an SS channel with the highest correlation (Gagnon et al., 2014).

For GLM, For HRF modeling, we fitted the GLM model to each subject. The GLM fitting here used two classes of regressors: HRF for each condition and the systemic signal clutter. The temporal basis functions used to model the HRF consisted of a sequence of 16 Gaussian functions, spaced 1 second apart, with a typical width of 1 second (Gagnon et al., 2011). This flexible model offers better fitting of the HRF shape at the expense of more parameter estimations than the typical canonical hemodynamic response function (Lindquist et al., 2009). Short-separation (SS) fNIRS signals were used as physiological regressors.

Fitting the GLM regression model to the entire data first before cross-validating the trials would result in information leakage (von Lühmann et al., 2020). In order to avoid leakage, we cross-validated both the GLM regression and classification steps. In each fold of the cross-validation, we fitted the GLM model to a training dataset and estimated regression coefficients using the Ordinary Least Squares (OLS) method. Then the short separation regression coefficients (SS coefficients) estimated from the training set are used for the test set where the individual trials are the difference between the measured fNIRS signals and the systemic physiological regressor weighted by the SS coefficients.

### Classification

As a baseline analysis, we used LDA with linear shrinkage of the covariance matrix (Ledoit and Wolf, 2004). We tested 2-class classification between left (−45°) and right (45°) spatial locations. The features used were the area under the curve of one-second-long segment of fNIRS signals. We performed 10 repetitions of 5-fold nested cross-validation (CV). Figure 2 shows the average CV accuracy across the subjects when all of the channels were used (from all targeted regions of interest).

**Figure 2:**
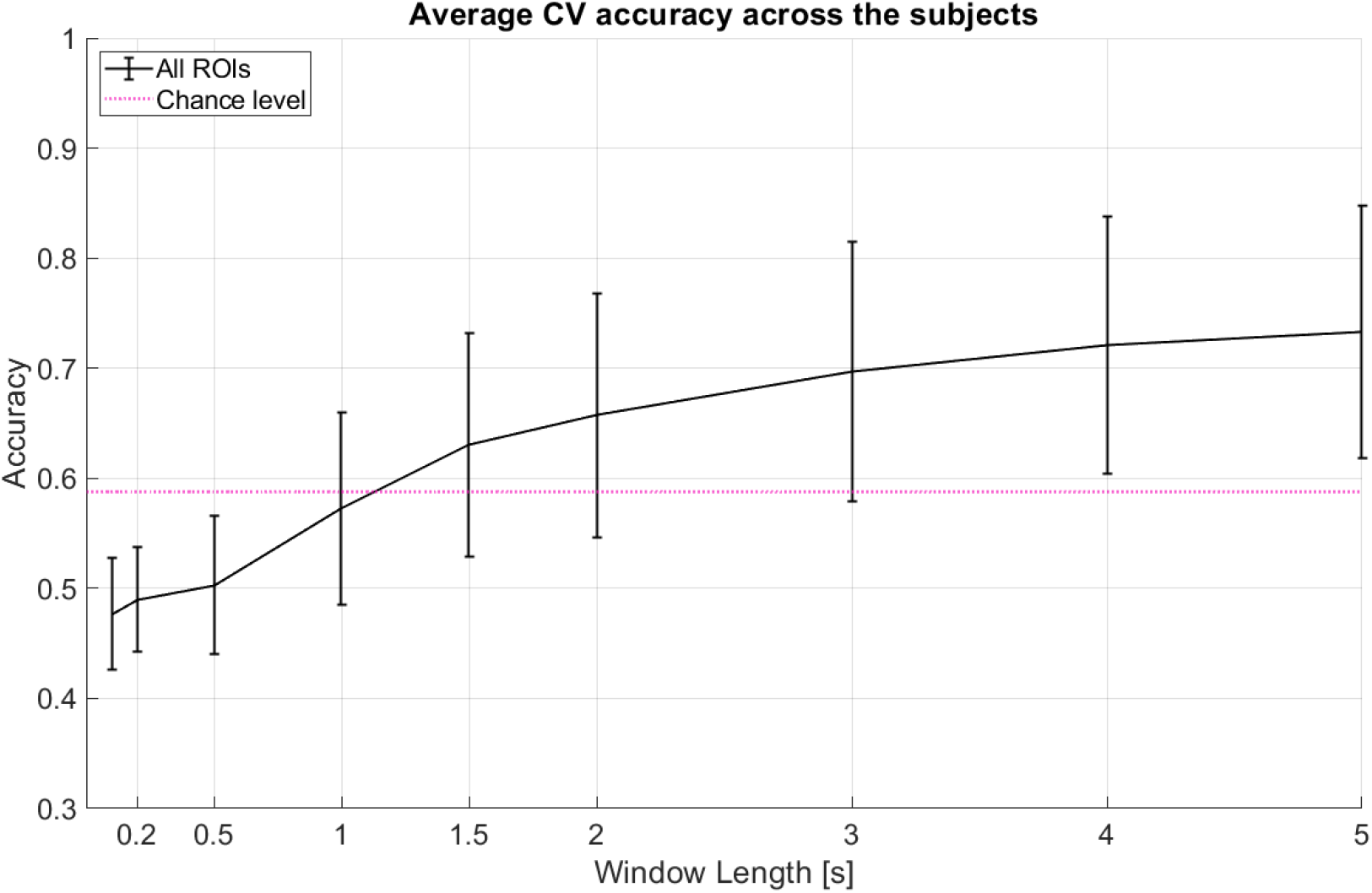
Average CV accuracy across 12 subjects when all regions of interest (ROI) are considered. All windows start at 0s (cue onset). Decision window lengths tested here are 0.1s, 0.2s, 0.5s, 1s, 1.5s, 2s, 3s, 4s, and 5s. The pink line indicates statistical significance for p = 0.05 using a two-tailed t-test and the given standard deviation from our cross-validation accuracies across subjects at window length 1s. Error bars represent 95% confidence intervals (counting each subject’s model performance accuracy as one sample point).

## SUMMARY

We presented the fNIRS dataset from 12 subjects, along with a behavioral dataset for decoding of attended audio-visual location. We include the custom MATLAB scripts for two-class classification (shown in baseline analysis) and three-class classification (not shown in figures). We also provide a link to the pre-processing code repository that includes any modification to the Master Homer3 package. We demonstrate via baseline analysis that our dataset can be used to train, test, and validate machine learning models for attention-based BCIs on a single trial basis. Recent and more advanced machine learning (such as deep learning neural network) models might give better accuracy results.

## DATA AVAILABILITY STATEMENT

All the data (fNIRS, probe design, and behavioral data) used in sample analysis in the paper are publicly available at doi:10.18112/openneuro.ds004830.v1.0.0. Matlab scripts used for analysis can be found in GitHub at https://github.com/NSNC-Lab/fNIRS_DecodingSpatialAttention.

## ETHICS STATEMENT

Recruitment of the subjects for this study was approved by Boston University IRB, and all of the IRB guidelines were followed. The subjects provided written consent for participation in the study, and they were compensated for their time.

## AUTHOR CONTRIBUTIONS

AL: methodology and writing – review and editing. DB: methodology and writing – review and editing. KS: conceptualization, project supervision and writing – review and editing. MN: methodology, experimental setup, data collection, algorithm and data analysis, writing – portions of original draft, writing – review and editing. MY: methodology, project supervision and writing – review and editing. SD: data analysis, data visualization, figure generation, writing – portions of original draft, writing -review and editing.

